# Anomaly detection for high-content image-based phenotypic cell profiling

**DOI:** 10.1101/2024.06.01.595856

**Authors:** Alon Shpigler, Naor Kolet, Shahar Golan, Erin Weisbart, Assaf Zaritsky

**Affiliations:** Department of Software and Information Systems Engineering, Ben-Gurion University of the Negev, Beer-Sheva 84105, Israel; Department of Computer Science, Jerusalem College of Technology, 91160 Jerusalem, Israel; Imaging Platform, Broad Institute of MIT and Harvard, Cambridge (MA), USA

## Abstract

High-content image-based phenotypic profiling combines automated microscopy and analysis to identify phenotypic alterations in cell morphology and provide insight into the cell’s physiological state. Classical representations of the phenotypic profile can not capture the full underlying complexity in cell organization, while recent weakly machine-learning based representation-learning methods are hard to biologically interpret. We used the abundance of control wells to learn the in-distribution of control experiments and use it to formulate a self-supervised reconstruction anomaly-based representation that encodes the intricate morphological inter-feature dependencies while preserving the representation interpretability. The performance of our anomaly-based representations was evaluated for downstream tasks with respect to two classical representations across four public Cell Painting datasets. Anomaly-based representations improved reproducibility, Mechanism of Action classification, and complemented classical representations. Unsupervised explainability of autoencoder-based anomalies identified specific inter-feature dependencies causing anomalies. The general concept of anomaly-based representations can be adapted to other applications in cell biology.

## Introduction

Visual cell phenotypes, characterized by the morphological features of a cell such as a cell shape and molecular composition, can serve as powerful readouts for cell state [1–6]. Alterations in visual cell phenotype, such as changes in cell shape and intracellular organization, can provide insight into the cell’s physiological state, as well as assist in the diagnosis and treatment of diseases [6–9]. High-content image-based phenotypic profiling combines automated microscopy and automated image analysis to identify phenotypic alterations in cell morphology [10–15]. For example, the Cell Painting assay uses high-content imaging, followed by analysis of multiple organelle stains in multiple cellular compartments at single cell resolution [16]. The most common approach for analysis of phenotypic profiling data, relies on pre-defined (a.k.a, “engineered”) single cell features, extracted with software tools such as CellProfiler [17], and followed by aggregation of these single cell features across the cell population to represent a “phenotypic profile”. The deviation of the treatment-induced phenotypic profile is measured in relation to the profile of untreated/control cells. The phenotypic alteration defines the degree and direction of change in the cellular high-dimensional phenotypic space, and can be used for various applications such as discovering drugs that “shift” a “disease-associated” to a “healthy-associated” phenotype or finding chemical compounds with a phenotype associated with a desired mechanism of action (MoA) [14, 18–20].

The gold standard for measuring a phenotype relies on the (inaccurate) implicit assumption of independence between the features extracted from the imaged cells. One example for this approach is measuring the fraction of features that dramatically deviate from the control profile in response to a treatment [14, 21]. Another example is measuring the correlations between profiles of treatments’ replicates [14, 22–23]. Of course, the profile features are interdependent, even after feature selection, because cell organization is so complex. For example, variation in cells’ intracellular organization may be largely explained by the cells’ shape [24]. Explicitly measuring these complex, non-linear dependencies for all features is not feasible due to the curse of dimensionality [25]. Thus, the biological function investigated may be incorrectly interpreted due to a simplified representation of the underlying data complexity. Recent representation-learning methods train machine learning models to encode lower-dimensional cell-level or well-level embeddings (called “*latent representations*”) through weakly supervised learning [26–31] or self supervised learning [32]. While representation-learning methods show promising results in terms of capturing the differences between treatments, current representation methods are not optimized to model the change posed by the treatment with respect to the control, but to distinguish between the different treatments (see Discussion). Moreover, representation-learning methods are harder to interpret, because the features forming the latent representations do not have a semantic cell biological explanation. Here, we propose using methods for anomaly detection to enhance the phenotypic profile representation in the context of high-content image-based phenotypic profiling by encoding intricate inter-feature dependencies while preserving the representation interpretability.

Anomaly detection aims at detecting abnormal observations that deviate from a predefined baseline pattern [33] and has vast applications in bioinformatics [34–35], healthcare [36], and cyber-security [37]. Anomaly detection relies on statistically characterizing the in-distribution of the data and defining observations that do not conform to this distribution as anomalous. Different approaches, such as neighbor-based [38–39] and isolation-based [40] were applied for anomaly detection, with ML emerging as an especially powerful technique in recent years [41]. The advantage of ML methods, and particularly deep neural networks, stems from their ability to integrate massive amounts of complex data into a generalized model that captures the inherent dependencies of the data distribution [42–43]. High-content image-based phenotypic profiling naturally aligns with the formulation of an anomaly detection problem with many replicates of the control condition that can be used to statistically define the in-distribution baseline patterns, and a few replicates from each of the (many) treatments, that can be assessed according to their deviation from the baseline, or in other words how anomalous they are in respect to the in-distribution baseline patterns.

We present here a reconstruction-based, self-supervised, anomaly detection-based representation for high-content image-based phenotypic cell profiling using the “gold-standard” CellProfiler representation. Reconstruction-based methods for anomaly detection are trained to reconstruct “normal” (i.e., non-anomalous) samples from low-dimensional encodings under the intuition that anomalous samples will be less successfully reconstructed due to altered dependencies between the representation’s features. We demonstrated that our anomaly-based representations derived from CellProfiler representations (hereafter called “anomaly-based” representations) surpass the raw CellProfiler representations (hereafter called “CellProfiler” representations) on multiple high-content Cell Painting datasets across different cell types and treatments [19], in terms of reproducibility and the downstream task of MoA identification. Moreover, we found that anomaly-based representations encapsulate complementary information in respect to the CellProfiler representations, leading to improved reproducibility and MoA identification. Finally, we demonstrated that applying an unsupervised method to explain autoencoder-based anomalies pinpointed specific inter-feature dependencies that changed to define an anomaly.

## Results

### Anomaly detection representation for image-based phenotypic cell profiling

Our anomaly detection-based method consists of three steps (Fig. 1). First, pre-processing, single cell feature extraction using CellProfiler [17] and well-level averaging across the cell population defining the well’s phenotypic profile (Fig. 1A). Second, leveraging the abundance of control wells, to train an “in-distribution” autoencoder deep neural network that encodes a lower dimensionality compressed phenotypic profile and decodes it to reconstruct the input profile in the original dimensionality, according to the (in-distribution) baseline of the control wells’ profiles (Fig. 1B). Half of the control wells from each experimental plate were pooled for training, and used to standardize all wells. Following feature selection performed on all profiles, the train controls are used for training the “in-distribution” autoencoder to minimize the discrepancy between the input and the reconstructed profiles. The trained autoencoder learns non-linear interrelationships between the features in the control profiles. Having access to all plates, the in-distribution autoencoder was guided to encode batch effects related to plate association, without the risk of exposing itself to data leakage, because the treated wells were not used for training. Third, the reconstruction errors of the in-distribution autoencoder are measured as the difference between the CellProfiler representations to the autoencoder output predictions, and define the anomaly-representations of treated wells (Fig. 1C). We estimated the in-distribution reconstructed error according to the control wells that were not included for training. The anomaly-representation of a treatment was defined as the deviation of the treated well’s reconstruction error, separately for each feature, in respect to the in-distribution controls’ reconstruction errors (Fig. 1D). High reconstruction errors indicate that the in-distribution autoencoder was not able to effectively reconstruct the wells’ profiles, suggesting that these treatments alter the morphological and intracellular organization in relation to control cells. These anomaly-representations can be used for downstream analyses such as hit identification in screening, assessment of treatment reproducibility and MoA identification [14, 19] (Fig. 1E). Importantly, the reconstruction error of each biologically-interpretable feature provides a direct “mechanistic” explanation, thus benefiting from transparent interpretability of hand-crafted features along with deep-learning capitalization of non-linearity and the wealth of control data.

**Figure 1.**
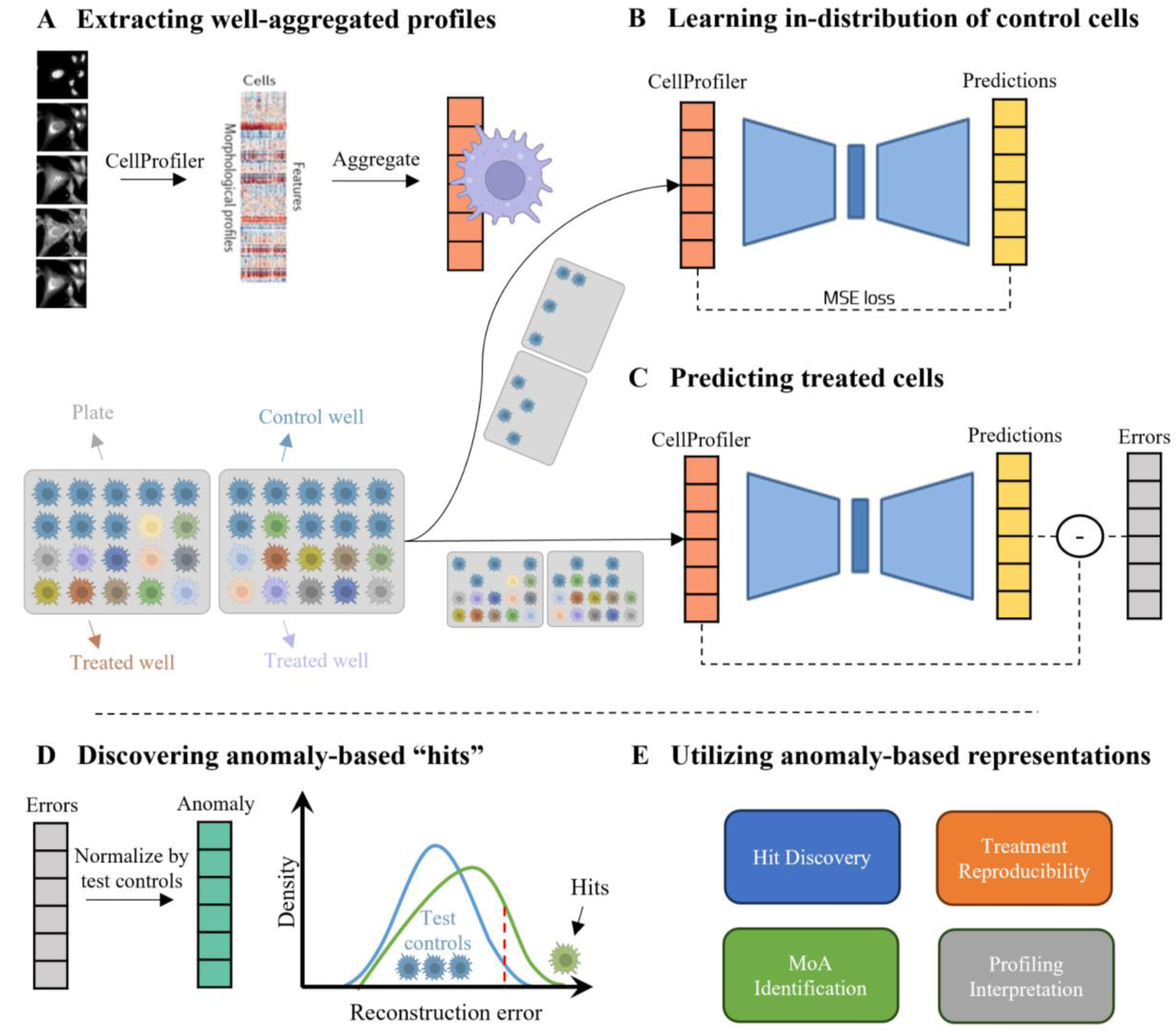
Method: anomaly-based representations for cell profiling. **(A) Top:** Well-aggregated profiles. Left-to-right: The CellProfiler software was used to extract single cell morphology features from Cell Painting images, the single cell features were aggregated to a well-level profile. **Bottom:** Each plate (gray) includes control (cyan) and treated (color) wells (cell icon). Half of the control wells were used to train the in-distribution autoencoder (B) and the other wells were used for evaluation (C). **(B)** The in-distribution autoencoder was trained to minimize the reconstruction error of control wells. **(C)** The in-distribution autoencoder was used to reconstruct control and treated wells. The reconstruction errors were calculated for each well. **(D)** The reconstruction error of a treated well was standardized according to the reconstruction errors of the control wells that were not used for training. These standardized reconstruction errors formed the “anomaly”-based representation. Treatments that lead to high reconstruction errors (green), in respect to the controls’ reconstruction errors (blue), were defined as “hits” (threshold in dashed red). **(E)** Anomaly-based representation can be used for a variety of downstream applications.

### Anomaly-based representations enhance reproducibility

Reproducibility, the extent to which a phenotype is replicated under the same experimental treatment, is a critical requirement in any screening application, where only a small fraction of treatments is followed-up with further comprehensive experimental validations (e.g., in drug discovery [45]). In high-content cell profiling, reproducibility serves as a measurement for the consistency of the experimental protocol and for the efficacy of the profile’s representation, and is used to exclude low-reproducible compounds from downstream analyses. We evaluated the reproducibility of our anomaly-representation in comparison to the CellProfiler representations in replicates of the same treatment across different plates. Reproducibility was measured by standardization of the well’s replicate-level profiles per plate, followed by measurement of the “Percent Replicating” score [14, 19, 22], the fraction of reproducible treatments, where a compound is deemed reproducible if its median pairwise profile correlation across replicates (“Replicate Correlation”), exceeds a threshold percentile of the pairwise correlation of random pairs of replicates across treatments (“Random Pairs Correlation”) (Fig. 2A-B). Following [19], we considered this threshold to be the 90th percentile of the random pairs’ distribution. We measured reproducibility of the anomaly- and of the CellProfiler-representations on four publicly available Cell Painting datasets from the Cell Painting Gallery [46] (Table 1): two compound screens (CDRP-bio [11], LINCS [14]) and two open reading frames (ORF) overexpression screens (LUAD [22], TAORF [44]); TAORF contains overexpression of WT cDNAs while LUAD contains overexpression of both WT and genetic variants. The median number of replicates per treatment (𝑁_𝑟_) in each dataset ranges between 5 to 8.

**Figure 2.**
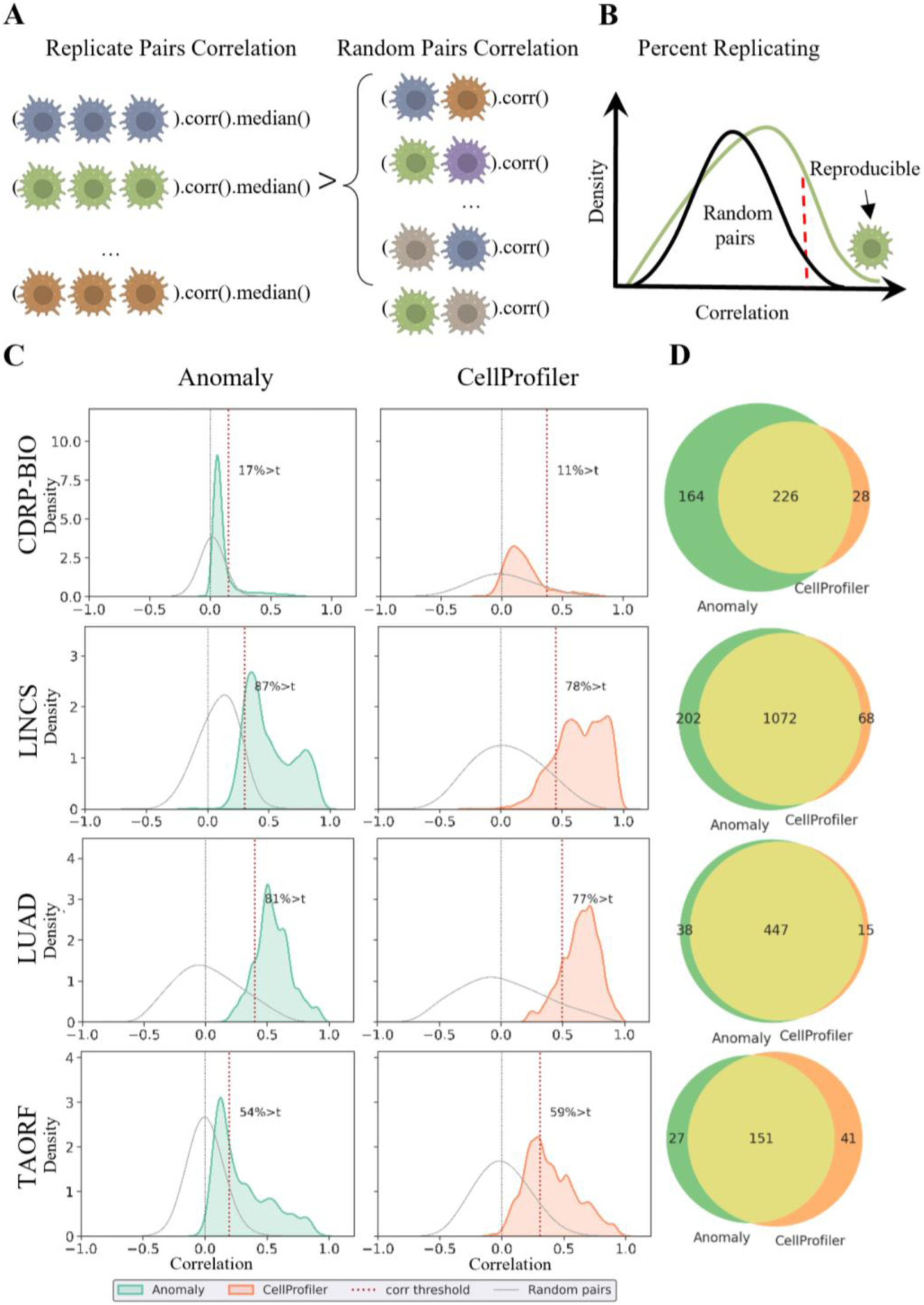
Anomaly-based representations are more reproducible. **(A)** Cartoon depicting the reproducibility determination. A treatment is defined as reproducible if the median pairwise correlation of its replicates (Replicate Correlation) is higher than the 90th percentile of the pairwise correlation among random replicates (Random Pairs). **(B)** Cartoon depicting the Percent Replicating score. The distribution of Replicate Correlations (green) versus the distribution of Random Pairs correlations (black). The dashed red vertical line defines the reproducibility threshold of 90% of the random pairs’ distribution - a well to the right of this line (green cell icon) is defined as reproducible, and the fraction of reproducible treatments determines the Percent Replicating score. (**C**) Percent Replicating scores across datasets for the anomaly-based (left) and the CellProfiler representations (right). Distribution of Replicate Correlations (green - anomaly-based, red - CellProfiler-based). Distribution of Random Pairs correlations (gray), zero correlation (dashed gray vertical line), reproducibility threshold (dashed red vertical line). **(D)** Venn diagram showing the number of reproducible treatments exclusive to the anomaly-based (green) or CellProfiler-based (orange) representations, and common to both representations (yellow).

**Table 1.** Datasets used in this study. Dataset: CDRP-bio (cpg0012-wawer-bioactivecompoundprofiling) [11], LINCS (cpg0004-lincs) [14], LUAD (cpg0031-caicedo-cmvip) [22], TAORF (cpg0017-rohban-pathways) [44]. Treatment: Chemical compounds, or ORF overexpression. Controls: number of control wells. Treatments: number of distinct treatments. 𝑁_𝑟_: median number of replicates per treatment.

| Dataset | Treatment Type | Cell line | Controls | Treatments | $N_r$ |
| --- | --- | --- | --- | --- | --- |
| CDRP-bio | Chemical | U2OS | 3,528 | 2,239 | 8 |
| LINCS | Chemical | A549 | 26,572 | 1,458 | 5 |
| LUAD | Genetic | A549 | 320 | 593 | 8 |
| TAORF | Genetic | U2OS | 120 | 324 | 5 |

The anomaly-based representations were more reproducible than the CellProfiler representations in three out of the four datasets (Fig. 2C and Table 2): percent replicating score of 17.41% versus 11.3% correspondingly (i.e., a 1.54-fold increase) in the CDRP-bio dataset, 87.38% versus 78.2% (i.e., a 1.12 fold increase) in the LINCS dataset, and 81.79% versus 77.91% (i.e., a 1.05 fold increase) in the LUAD dataset. For TAORF, the smallest of the inspected datasets, CellProfiler representations reproduced 59.26% of the ORF overexpression versus 54.94% reproduced by the anomaly-based representation (i.e., a 1.08-fold increase in favor of CellProfiler). The deterioration in reproducibility could be related to the limited number of control wells in the TAORF dataset, with only 60 (out of 120) control wells used for training the in-distribution autoencoder. For comparison, the CDRP-bio dataset has 3,528 control wells (Table 1). Systematic assessment of the percent replicating score as a function of the number of control wells verified that increased numbers of control wells enhanced the anomaly-based representations reproducibility, probably by learning a more accurate representation of control well’s in-distribution (Supplementary Fig. 1). Low performance was observed for TAORF and CDRP-bio datasets on previous research and was hypothesized to derive from poorer technical data quality [19]. Together with the low number of training samples, that may lead to inaccurate in-distribution learning. The distributions of Replicate Correlations were lower for the anomaly-based representation, regardless of being more reproducible in comparison to CellProfiler representations, because their corresponding Random Pairs Correlations were more concentrated around 0 (Fig. 2C). These low correlations among random replicates suggested that the anomaly-based representations were less sensitive to experimental batch effects that may lead to spurious correlations. We next evaluated what fraction of the reproducible treatments was common and what fraction was distinct for the anomaly-based and CellProfiler representations (Fig. 2D). Most of the reproducible treatments were consistently identified in both representations, with larger numbers of reproducible treatments exclusively found by the anomaly-based representation for all datasets excluding TAORF. The union of the reproducible treatments found by either representation leads to a major improvement in the numbers of reproducible treatments that can be used for downstream analyses such as identifying their Mechanism of Action: 64% more in CDRP-bio, 17% in LINCS, 8% in LUAD, and 14% in TAORF. Finally, we evaluated whether the anomaly-based representations also provided complementary information to the mRNA expression of 978 genes (called the *L1000* profile). Analysis of the complementary of all three representations showed that the anomaly-based representations are complementary to both CellProfiler and L1000 (Supplementary Fig. 2). Altogether, these results indicate that anomaly-based representations encapsulate complementary information with respect to the CellProfiler representations that can lead to the availability of more reproducible treatments for follow-up investigations.

**Table 2.** Reproducibility results. Percent Replicating score for the CellProfiler (left column) versus the anomaly-based (middle column) representations. The union of treatments found reproducible by either representation is shown in the right column (𝐴 ∪ 𝐶).

| Dataset | CellProfiler (%) | Anomaly (%) | A ∪ C (%) |
| --- | --- | --- | --- |
| CDRP-bio | 254 (11.3) | 390 (17.41) | 418 (18.66) |
| LINCS | 1140 (78.2) | 1274 (87.38) | 1342 (92.04) |
| LUAD | 462 (77.91) | 485 (81.79) | 500 (84.31) |
| TAORF | 192 (59.26) | 178 (54.94) | 219 (67.59) |

### Anomaly-based representations enhance Mechanism of Action identification

Linking a compound to its mechanism of action (MoA) is an important application of image-based phenotypic profiling [14, 18–20, 47]. We evaluated the ability to identify the MoA of compounds using anomaly-based representations in comparison to the CellProfiler representations. CDRP-bio and LINCS include compounds with known MoA and were thus used for this evaluation. We followed a previously described workflow to benchmark MoA identification on these datasets [14, 19] (Fig. 3A). First, we included compounds that were found reproducible by one of the anomaly-based, CellProfiler, or L1000 representations. Second, we excluded MoAs with fewer than 5 reproducible compounds linked to them. The complementarity between anomaly-based, CellProfiler and L1000 representations leads to inclusion of more compounds that leads to inclusion of more MoAs with sufficient numbers of treatments. Third, we evaluated compounds’ MoAs predictions for different representations, with a 5-fold cross-validation using Logistic Regression (LR) and Multi Layer Perceptron (MLP) classifiers. This process was repeated 10 times for robustness analysis. The comparison between the anomaly-based and the CellProfiler-based representations was performed using two inclusion criteria for selecting reproducible compounds to train MoA classifiers: according to (1) CellProfiler and L1000 (𝐶 ∪ 𝐿), or according to (2) Anomaly, CellProfiler, and L1000 (𝐴 ∪ 𝐶 ∪ 𝐿). Inclusion of reproducible compounds determined by the anomaly-based representations (𝐴 ∪ 𝐶 ∪ 𝐿) increased the number of MoAs (38% more for CDRP-bio and 12% more for LINCS), and increased the total number of reproducible compounds (36% for CDRP-bio and 18% for LINCS) (Fig. 3B). For each of the two inclusion criteria subsets the anomaly-based representations exhibited superior performance compared to the CellProfiler baseline for both the LR and the MLP classification models, with an average weighted F1-score of 0.301 versus 0.238 for the best model trained with anomaly-based versus CellProfiler representations correspondingly for the CDRP-biodataset, and 0.212 versus 0.183 for the LINCS dataset (Fig. 3C). Specifically, the anomaly-based representations lead to prominent improvements in the CDRP-bio mechanisms of actions CDK inhibitor (ΔF1-score = 0.305), adrenergic receptor agonist (ΔF1-score = 0.167) and p38 MAPK inhibitor (ΔF1-score = 0.165), and in the LINCS MoAs for vitamin D receptor agonist (ΔF1-score = 0.207), EGFR inhibitor (ΔF1-score = 0.138), and in gamma secretase inhibitor (ΔF1-score = 0.125) (Fig. 3D). Additionally, anomaly-based representations were also found to be complementary to the genetic L1000 profiles, showing improved results when concatenated together compared to using each one independently (Supplementary Fig. 3). Altogether, anomaly-based representations improve MoA classification by both increasing the number of reproducible compounds available for MoA classification and by encoding more discriminative information than the CellProfiler-based representations.

**Figure 3.**
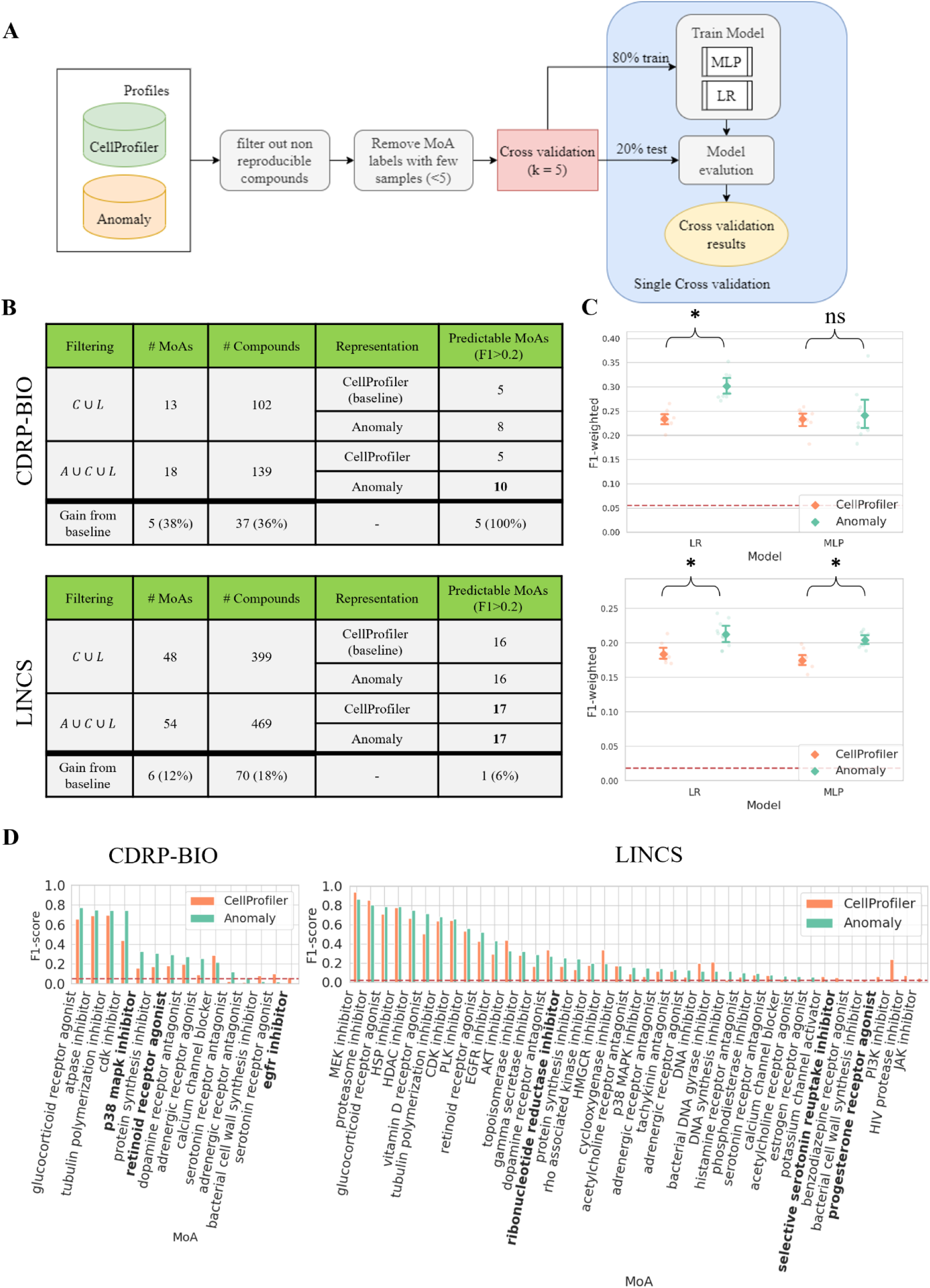
Anomaly-based representations improve MoA classification. (**A**) MoA classification workflow. Left-to-right: inclusion of compounds that met our reproducibility criteria (for either the CellProfiler or the Anomaly-based representations), followed by exclusion of MoAs with < 5 compounds attributed to them. Training machine learning models using the anomaly-based versus the CellProfiler-based representations with cross-validation. (**B**) MoA classification results for anomaly-based versus CellProfiler-based representations, using the reproducible inclusion criteria with (𝐴 ∪ 𝐶 ∪ 𝐿) or without (𝐶 ∪ 𝐿) the anomaly-based representations. (**C**) F1-scores of anomaly-based (green) versus the CellProfiler-based (orange) representations trained on MLP and LR models for CDRP-bio (cpg0012-wawer-bioactivecompoundprofiling) and LINCS (cpg0004-lincs) datasets. Each dot represents an experiment and bars indicate confidence intervals. The red dashed line indicates the F1 random score. * - statistically significant (p-value < 0.05) by Welsh t-test, ns - not significant. P-values for CDRP-bio were 5𝑒^−6^ (LR) and 0.66 (MLP), and for LINCS 0.001 (LR) and 3𝑒^−5^ (MLP). **(D)** MoA-specific F1 scores using the (better performing) LR model. Bold indicates MoAs that would not be included without reproducible compounds according to the anomaly-based representations. F1-scores lower than the random score for both representations (3/18 MoAs in CDRP-bio, and 12/54 MoAs in LINCS) were excluded from the figure to improve clarity.

### Anomaly-based representations are interpretable

Understanding the underlying reasons behind a treatment’s hit identification is crucial for meaningful interpretation and subsequent decision-making. The CellProfiler hand-crafted features are linked to specific morphological phenotypes, and thus treatment-induced alterations of many of these features can be biologically interpreted and investigated. Interpretation of the anomalous features is more challenging because an anomalous (i.e., poorly reconstructed) feature can be “caused” by a combination of subtle alterations in several inter-dependent input features (Fig. 4A). To interpret our anomaly-based representation we applied a recent unsupervised extension of the widely used SHAP (SHapley Additive exPlanations) [48] that was designed to explain anomalies identified by an autoencoder [49]. SHAP assigns importance values to features based on their additive contribution to the model output, and its extension to autoencoder-based anomalies isolates the input features contributing to the high reconstruction errors by treating each feature as a separate reconstruction task (Fig. 4A).

**Figure 4.**
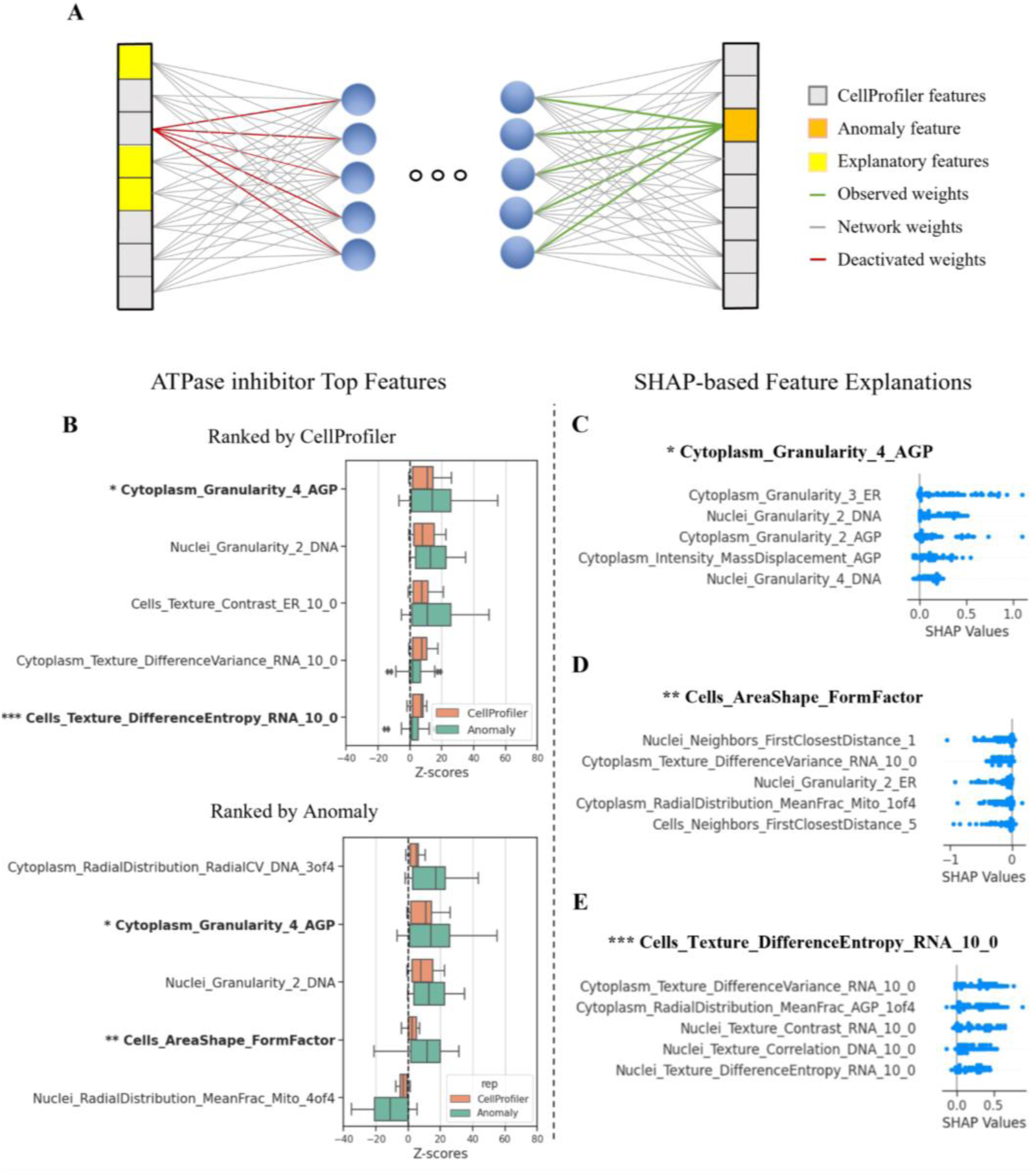
MoA interpretation. (**A**) Illustration of the explanation process of the anomalous features (orange on the right). This anomaly is explained by the combined alteration of multiple input features (yellow on the left). A group of well-level profiles of interest are passed through the network, and the features exhibiting the highest reconstruction errors are identified. For each investigated feature (orange square in output), the weights leading to its activation (green lines) are activated and the weights going from its CellProfiler representation in the input are deactivated (red lines). The well-level profiles are reintroduced into the network and the features that led to high reconstruction errors are found by using the autoencoder-based anomaly SHAP. These explanations can be pooled according to a treatment or MoA to provide the corresponding explanation. (**B**) Distributions of ATPase inhibitor top five features’ z-scores for the CellProfiler (orange) and anomaly-based representations (green) ranked according to the CellProfiler median z-scores (top) and by anomaly median z-scores (bottom) in the CDRP-bio dataset.Features in bold are analyzed using the autoencoder-based anomaly SHAP in the following panels. **(C-E)** Explanations for selected (see text for justification) altered features in the ATPase inhibitor MoA in the CDRP-bio dataset. Each dot represents a replicate well, and the x-axis represents the SHAP values contributing to the errors. Negative values indicate an inverse relationship between the inspected feature and the input feature compared to the relationship in the control population.

To illustrate how SHAP can be used, we focused on the interpretation of the ATPase inhibitor MoA, which was one of the top predicted MoA classes in the CDRP-bio dataset (Fig. 3D). We first examined a feature that was high in both CellProfiler and anomaly representations. The CellProfiler feature “Cytoplasm_Granularity_AGP_4” was ranked first in terms of its deviation from the control, and was ranked second in the anomaly-based representation (Fig. 4B, single asterisk). Granularity measures the signal lost with iterative erosions relative to the total signal and can be hard to directly interpret. The autoencoder-based anomaly SHAP explanation for “Cytoplasm_Granularity_AGP_4” included the “adjacent” Granularity feature, “Cytoplasm_Granularity_AGP_2”. This result aligned with a recent study [50] reporting that genetically disrupting the Vacuolar ATPase (which is a specific ATPase) caused a specific decrease in the first Granularity feature in the Golgi and plasma membrane channel (the most similar channel to the F-**a**ctin, **G**olgi, and **p**lasma membrane (AGP) channel in the CDRP-bio dataset), and a concomitant increase in other Granularity features. Our feature selection excluded some AGP Granularity features so we retrained the in-distribution autoencoder using the full feature set and were able to observe a similar phenotype for ATPase inhibitor MoA in both the CellProfiler and the anomaly-based representations (Supplementary Fig. 4). Though none of the compounds in the ATPase inhibitor MoA class in CDRP-bio was annotated to directly target the Vacuolar ATPase, the similarity of our observations can easily be explained by either perturbation causing a general disruption of intracellular acidification and therefore similar observable phenotypes.

A feature with a high reconstruction error but an unperturbed CellProfiler value implies that the in-distribution autoencoder reconstruction was hampered by the combined effect of alteration in other features. An example is “Cells_AreaShape_FormFactor” (Fig. 4B, double asterisk). The FormFactor feature measures the roundness of objects by calculating the ratio between the object area and perimeter, i.e., larger values indicate more circular objects. The autoencoder-based anomaly SHAP explanation of this feature included two features that measure the cell’s distance from neighboring cells (“Neighbors_FirstClosestDistance” features), with higher distances in the input leading to lower cell roundness in the reconstruction (Fig. 4D). Typically, distance from neighbors is a measure of local cell sparsity, and cells at sparse environments have more space to spread and thus may have lower FormFactor values. Intriguingly, this relationship is affected in this MoA and further exploration could be done to test how frequently this relationship is broken across MoA classes and whether this is correlated with compound toxicity as AreaShape features are known to play an important role in predicting Cell Health readouts [51].

A feature with a low reconstruction error but a perturbed CellProfiler value implies that the inter-relations learned in the in-distribution autoencoder were maintained and thus the deviation of the feature can be properly reconstructed by using alterations of other features. An example is “Cells_Texture_DifferenceEntropy_RNA_10_0” (Fig. 4B, triple asterisk). This feature’s autoencoder-based anomaly SHAP explanation identified several positive relations with other texture features (Fig. 4E). As texture features are highly represented in Cell Painting datasets but are not human-interpretable, in this example a researcher could instead focus their follow-up experiments on the interpretable radial distribution feature of AGP in the cytoplasm as a welcome alternative approach to understanding the biology behind low interpretable CellProfiler features.

Cumulatively, our results indicate that autoencoder-based anomaly SHAP explanation of anomaly-based representations can reveal specific biological-interpretable dependencies between features that break following a treatment.

## Discussion

We take advantage of the abundance of control wells, to formulate the high-content image-based cell profiling as an anomaly-detection problem. Our anomaly-based representations learn the inter-feature complex dependencies of the population of “normal” (i.e., unperturbed) cells by minimizing the control reconstruction errors under the premise that perturbations in the cells’ organization will lead to less successful reconstruction due to altered inter-feature dependencies. Such representations hit a sweet-spot in terms of the inherent tradeoff between performance and interpretability. First, our anomaly-based representations model the dependencies between features, surpassing traditional methods of analyzing CellProfiler-representations where features are considered independently in reproducibility (Fig. 2) and MoA classification (Fig. 3). The low correlations among random replicates (Fig. 2C) suggests that these representations implicitly mitigate batch effects, a known confounder of cell profiling, meriting the future application of anomaly-based representations for batch correction [52]. Second, optimizing representations to capture these inter-feature dependencies provide a complementary readout that can identify perturbations that extensively alter these dependencies, without necessarily sufficiently altering each of the individual features (Figs. 2-3). For example, several positively correlated features may lead to an anomaly due to a marginal increase of some features along with a marginal decrease of other features. These subtle phenotypes are not sufficiently profound to create a reproducible phenotype, without encoding the deviation of the non-linear convoluted dependencies. Third, using the CellProfiler-derived features as the starting point for our anomaly-representations make the latter more interpretable with respect to most deep learning “black-box” representations. Specifically, using unsupervised explainability we were able to extract biologically meaningful dependencies that deviated in response to a perturbation (Fig. 4).

Weakly supervised representation-learning uses experimental labels, such as the treatment label, to “guide” their representations such that replicates of the same label are encoded close to one another in the latent space and different labels are encoded far apart [26–31]. Beyond optimizing representations that are hard to biologically-interpret, experiment treatment-based weak supervision may lead to undesired consequences where the representations of treatments that lead to a similar phenotype are pushed away from one another in the latent space because they do not share the same label, possibly even pushing one representation closer to the control (Supplementary Figure 4A). In turn, representations with reduced cross-treatment phenotypic similarity may induce errors in downstream analyses, especially in unsupervised interpretation of biological function such as lead optimization [53] and identifying unknown MoA signatures [14,19]. In contrast, our anomaly-based method is self-supervised, i.e., does not use treatment labels (or any other assumptions on the underlying data) to guide the representation, rather treatment profiles’ latent representations are encoded solely according to their deviation from the control in-distribution (Supplementary Fig. 5B).

The core idea of learning the in-distribution of control experiments followed by identifying anomalies in respect to this distribution is a general concept that can be adapted to applications in cell biology beyond high-content image-based cell profiling. For example, recent work suggested a form of anomaly detection to identify treatments leading to altered cell fate (apoptosis or mitosis) decision processes during live imaged cell competition assay by training a model for the prediction of “normal” cell fate behavior and a discriminator network to determine anomalies deviating from the expected cell decision process [54]. Of course, any anomaly-based representation requires sufficient control replicates to properly model the in-distribution of the data (see Supplementary Figure 1). In applications where there are no controls, but sufficient data, anomaly-based representations can still be applied by learning the in-distribution of one experimental condition and then modeling the reconstruction errors of the other experimental conditions in respect to the in-distribution reconstruction errors. For example, in single cell spatial multiplexed proteomics applied to clinical data, each patient has many cells, and thus we can learn the in-distribution of the protein expression across cells from one disease state and apply it to patients from other disease states. Our application involved first deriving pre-defined features from the raw image and then applying anomaly detection on the tabular representation. A similar approach can be applied directly to the raw image data using convolutional neural networks (e.g., [55]). Learning the in-distribution directly from the raw images avoids segmentation errors and can unbiasedly encode spatial relations that were not measured by the CellProfiler features, but suffer from poor interpretability. Imaged-based anomaly detection can also have applications in other domains such as single cell spatial omics.

## Methods

### Datasets

All datasets were created at the Broad Institute and previously published. Two datasets had chemical perturbations and two had genetic perturbations (Table 1). For each dataset two distinct readouts were recorded: gene expression (GE) profiles and morphological profiles (Cell Painting). Each dataset was acquired by plating cells in two sets of identical plates, perturbed identically in the same laboratory. One of these sets was used to measure GE and the other set was imaged to measure morphology. The morphological profiles were captured using the Cell Painting assay [16]. Cell painting is a microscopy-based assay that acquires five fluorescence channels labeling of the actin cytoskeleton, Golgi apparatus, plasma membrane, nucleus, endoplasmic reticulum, mitochondria, nucleoli and cytoplasmic RNA. Acquired microscopy images were processed using the CellProfiler software [17] to extract 1,783 features of each cell’s morphology such as shape, intensity and texture statistics, that were aggregated (population-averaged) to create per-well profiles. The features are measured in three cell regions – nucleus, whole-cell, and cytoplasm (difference between whole-cell area and nuclei area) . The GE profiles were acquired using the L1000 assay [56], that provides high-throughput measurement of mRNA levels for 978 genes, roughly covering 82% of the transcriptional variance across the entire genome. Cell Painting profiles were used for anomaly detection representations. GE profiles were used as additional input for selecting reproducible treatments and for MoA classification. We downloaded all metadata-augmented per-well aggregated Cell Painting datasets from the Cell Painting Gallery (CPG) (https://registry.opendata.aws/cellpainting-gallery/) [46]. We downloaded the preprocessed profiles and performed normalization and feature selection in the training process to avoid data leakage during normalization (See Data preprocessing section) (See Data preprocessing section). GE profiles were also downloaded from the CPG [46].

### Data split

Each experimental plate had multiple control replicates and many treated wells without per-well replicates (i.e., one replicate per treatment in a well). Per plate, the control wells were randomly split to 40%:10%:50% between train, validation and test, correspondingly. Thus, each plate’s wells were split to four subgroups of wells: (1) training controls, (2) validation controls, (3) test controls, and (4) treated wells for evaluation at inference.

### Data preprocessing

Feature-wise standardization was implemented on all well-level samples using the aggregated training controls population. Normalization was exclusively conducted on the training set to prevent any leakage of the test data distribution during the normalization process. Whole-plate normalization was performed after training (See “Measuring reproducibility” section). Feature selection was performed using the Pycytominer software [57], removing features with high ratio of null values (null_threshold = 0.05), high correlation (corr_threshold=0.9), and low variance (freq_cut = 0.05, unique_cut = 0.01), all with the default parameters as set by Pycytominer. Following feature selection, the number of features dropped from 1,700 to a range of 300-700 for all datasets.

### Anomaly detection

The autoencoder architecture consisted of a large encoder and a flat decoder. Three-layer encoder, with layer sizes 256-128-64, where each layer contains a Relu activation layer. The latent dimension was of size 32. A 1-layer decoder transformed the latent representation back to the input size. The loss function was the mean square error between the reconstructed output and the corresponding observed CellProfiler representation, with an additional l2 regularization loss to restrict the output amplitude. Each autoencoder (one per dataset) was trained until convergence or a maximum of 300 epochs with batch size of 32. Hyperparameter optimization was applied for each dataset with a learning rate range of 1e-5 to 1e-3, output features l2 regularization range of 1e-5 to 1e-2, and dropout ratio of 0 to 0.2. Total time for training each of the datasets, including hyperparameter optimization, was up to one hour on the Nvidia RTX1080 GPU.

At inference, the trained autoencoders calculated the feature-wise predictions’ reconstruction error for each well called the anomaly-based representation. All treatment replicates were z-score normalized according to the test control wells. These steps resulted in a vector of feature-wise z-scores for each treatment replicate.

### Measuring Reproducibility

Whole-plate normalization was applied based on all replicate-level z-scores, including the control and the treatment replicates, because it was the optimal setting for reproducibility, in agreement with previous studies [19, 50]. For all representations (CellProfiler-based, Anomaly-based, L1000), reproducibility was measured according to the fraction of reproducible treatments, a measurement called the “Percent Replicating” score [14, 19, 22]. Briefly, for each treatment, we first calculated the mean Pearson correlation coefficient of each pair of replicates profiles. Second, we compared the median replicate pairs correlation of each treatment to a null distribution containing correlations of all random pairs of treatments. Third, for robustness, we repeated the previous step for five times, altogether generating a null distribution containing 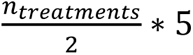 samples. Last, following [19], we defined a treatment as reproducible if its mean Replicate Correlation was above the 90^𝑡ℎ^ percentile of the null distribution. The LINCS had multiple doses per compound, and reproducibility was measured based on replicates of the highest dose for each compound.

### MoA classification

We compared MoA classification based on CellProfiler-based and anomaly-based representations for the CDRP-bio and LINCS datasets that included compounds with known MoA (Supplementary Fig. 6). First, for each representation, each treatment was represented by its mean profile across plate-normalized replicates. Second, following [19], we selected reproducible treatments according to two inclusion criteria: (1) Reproducible treatments according to the CellProfiler or L1000 representations, or (2) Reproducibility according to the Anomaly, CellProfiler, or L1000 representations. Third, we excluded all MoAs with fewer than five reproducible treatments. All treatments’ features were scaled to a range of [0,1] and were used to train a Logistic Regression (LR) and a Multi Layer Perceptron (MLP) MoA classifiers.

Each model was applied for prediction of MoA labels using CellProfiler-based and anomaly-based representations independently to compare the representations. We performed stratified nested k-fold cross-validation (k = 5) to evaluate the classification performance. The MLP classifier architecture was made of two layers with a Relu activation layer between them. Hidden layer sizes, regularization strength, and learning rates were optimized per training fold. LR was optimized with different regularization parameters per training fold. MoAs vary in compound numbers, ranging from five to 25 (Supplementary Fig. 6). To address data imbalance, models underwent oversampling to align class sizes with the majority class in the training set. The predictions were evaluated using the F1-score metric. A naive random F1 score for classes was calculated for baseline. Each classification experiment was repeated 10 times for robustness. The statistical significance of the difference between CellProfiler and Anomaly-based representations was calculated using the Welsh t-test, a two-sample test used to test if two populations have different means. The test was performed with N=10, the number of experiments per setting.

### Autoencoder-based anomaly SHAP explanations

To interpret what were the inter-feature associations that deviated following a treatment we used an extended version of the classic Shapley additive explanation (SHAP) method [48] that was designed to explain anomalies identified by an autoencoder [49]. For a treatment or MoA of interest, the autoencoder-based anomaly SHAP explanations were obtained independently for each anomalous feature by calculating the autoencoder reconstructed output values of that feature and then the SHAP values of the input features in relation to the output feature at test.

First, the most anomalous features were identified according to the reconstruction errors. Second, given an anomalous feature, the weights associated with this feature in the first layer were deactivated and the well-level replicate profiles were re-introduced to the network. The output values of the inspected feature were analyzed using kernel SHAP to compute the SHAP values of the input features, i.e. the importance of each input feature in predicting the anomalous feature. Kernel SHAP is a model-agnostic approximation of SHAP values via weighted linear regression. The Kernel SHAP estimates the additive value of features in the input space for a subset of samples of interest. The importance of an input feature is determined by the change in the model’s output when that input feature is set to “missing”. The “missing” values are replaced with values taken from a reference dataset that reflects the distribution of the general data. We used samples from the control test set as the reference dataset for kernel SHAP approximation.

### Source code and data availability

Data used for this study was downloaded from the Cell Painting Gallery (CPG) (https://registry.opendata.aws/cellpainting-gallery/) [46]. Specifically, all datasets were collectively downloaded from cpg0003-rosetta. This study did not generate new unique datasets. Source code is publicly available at https://github.com/zaritskylab/AnomalyDetectionScreening.

## Supplementary Material

**Supplementary Figure 1.**
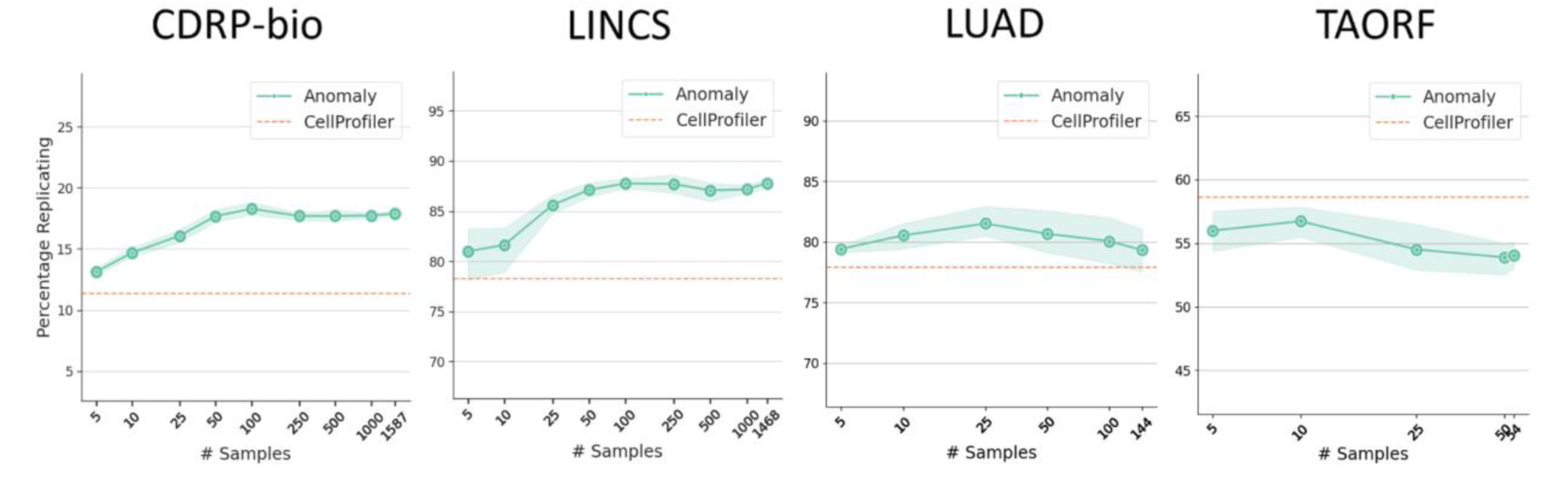
Percent replicating score as a function of the number of control wells used to train the in-distribution autoencoder. Dashed orange line - the percent replicating score of the CellProfiler representations. Green data points and shade show the percent replicating score mean and the standard deviation of the anomaly-based representations over 5 independent experiments.

**Supplementary Figure 2.**
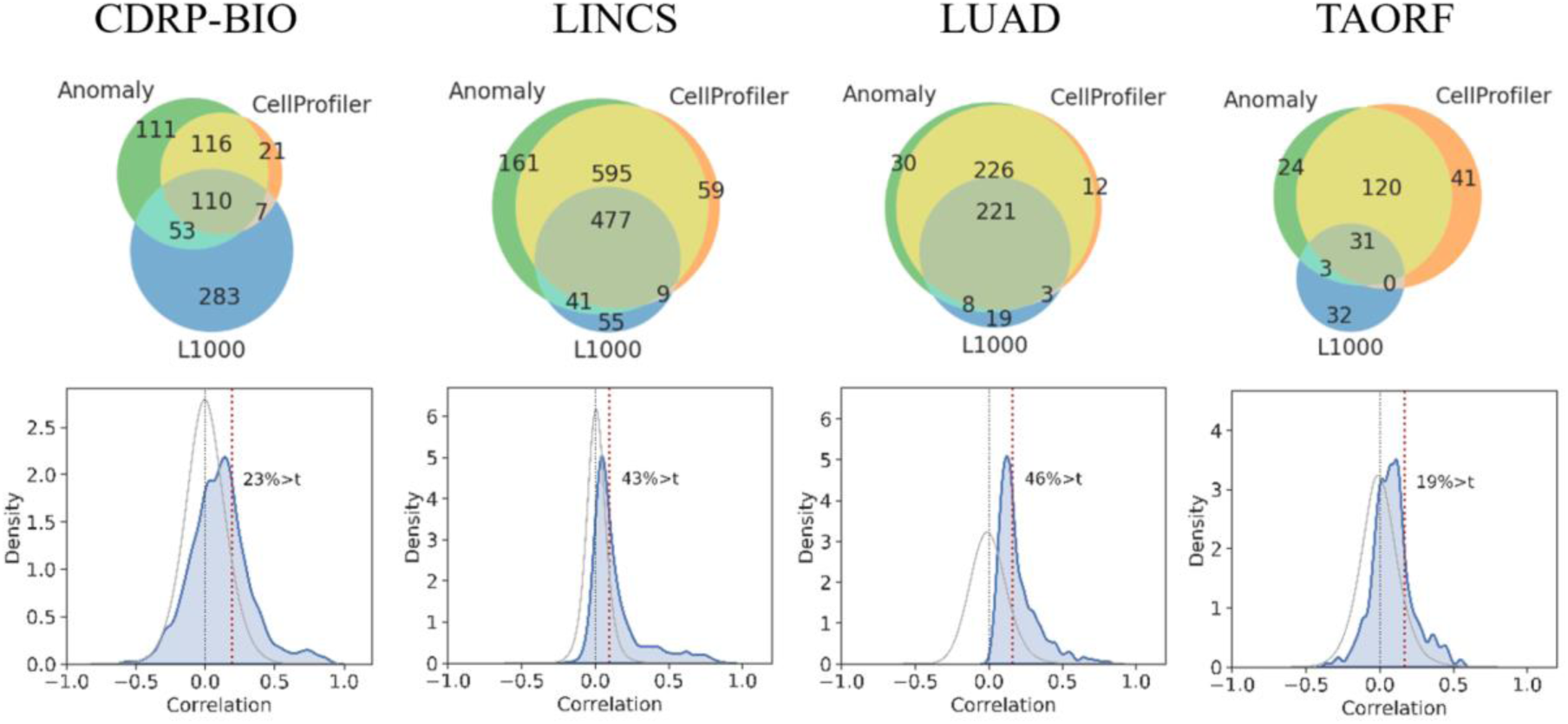
Analysis of the complementary information shared between the anomaly-based, the CellProfiler, and the gene expression representations (L1000). **Top:** Venn diagram showing the number of reproducible treatments exclusive to the anomaly-based (green), CellProfiler-based (orange), and L1000-based (blue) representations. Treatments found reproducible by multiple representations are shown by the intersection of the different circles. **Below**: Percent Replicating scores for the L1000-based representations. Distribution of Replicate Correlations (blue), Random Pairs Correlations (gray), reproducibility threshold (dashed red vertical line).

**Supplementary Figure 3.**
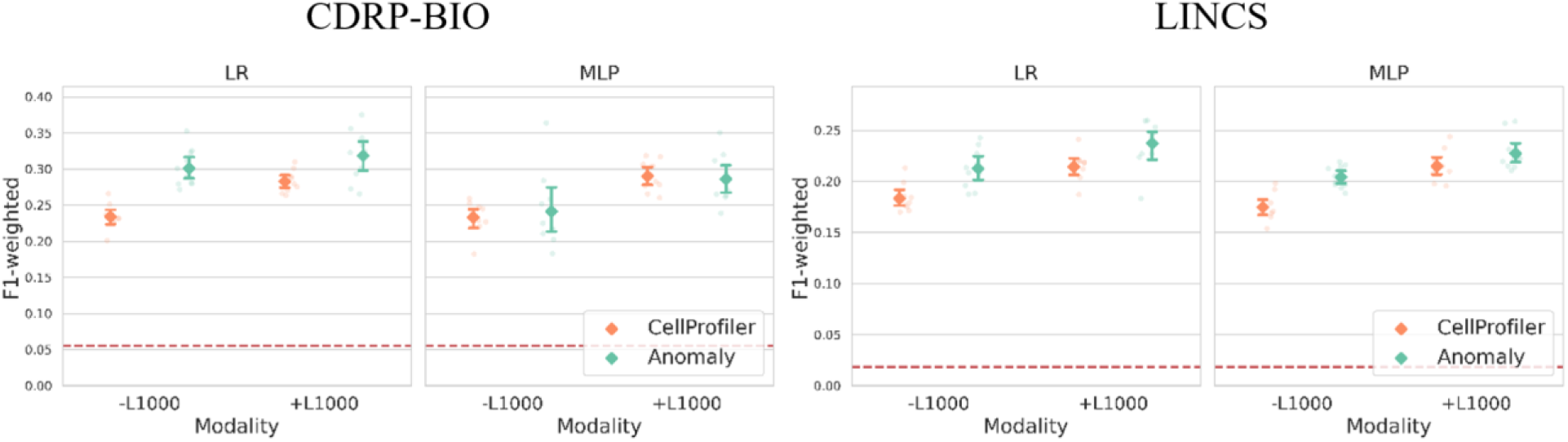
Genetic expression representations (L1000) complement the CellProfiler (orange) and anomaly-based (green) representations for MoA classification. Logistic Regression (LR) and Multi Layer Perceptron (MLP) MoA classification F1 performance on CellProfiler versus anomaly-based representations without (-L1000) or with (+L1000) the concatenation of the L1000 representation. Each dot represents a unique experiment, and bars indicate 95% confidence intervals. Red dashed line indicates the F1 random score.

**Supplementary Figure 4.**
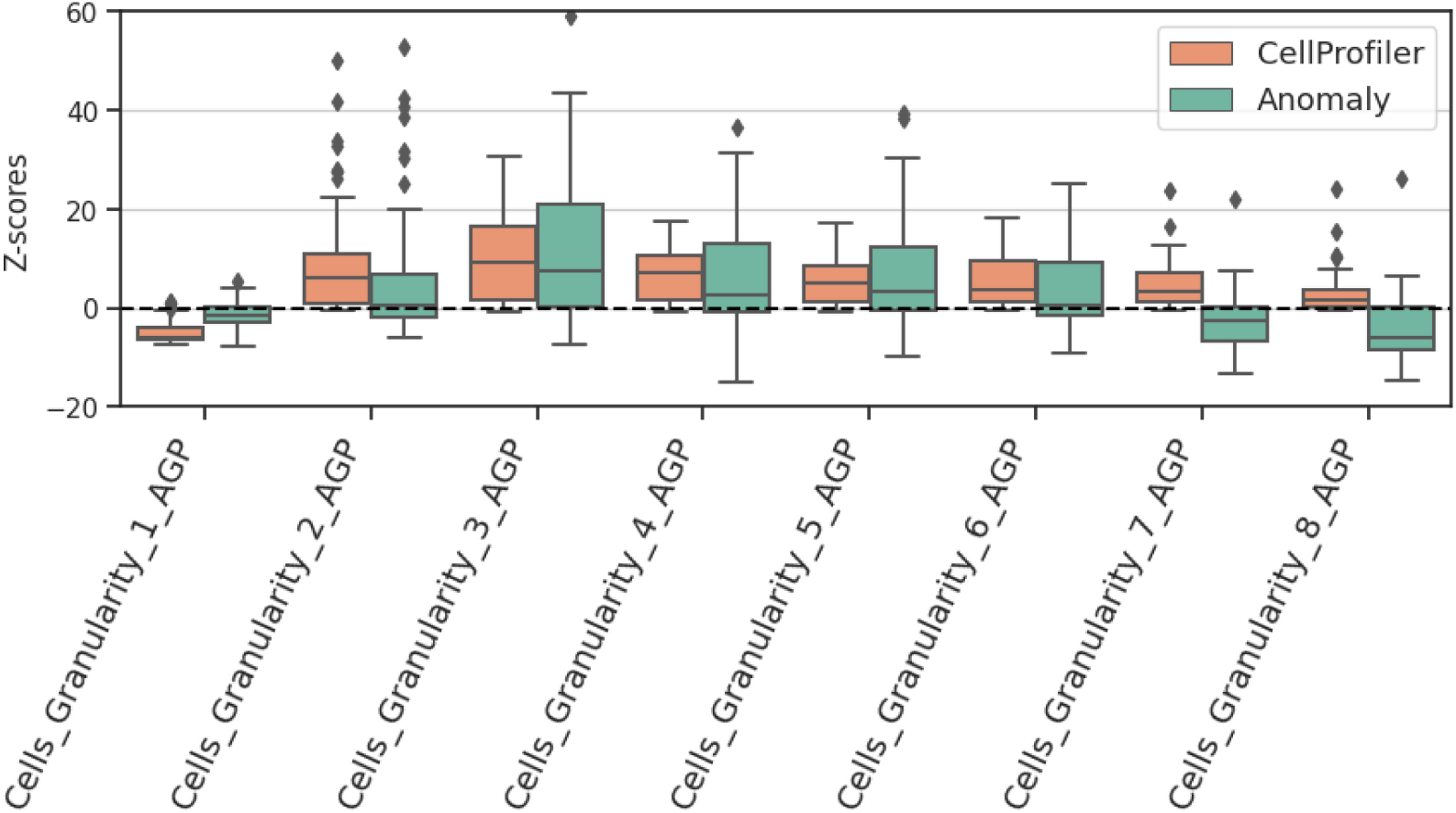
Granularity analysis in the AGP channel for the MoA of ATPase inhibitor in the CDRP-bio dataset. Feature z-scores distributions for CellProfiler (orange) and anomaly-based (green) representations.

**Supplementary Figure 5.**
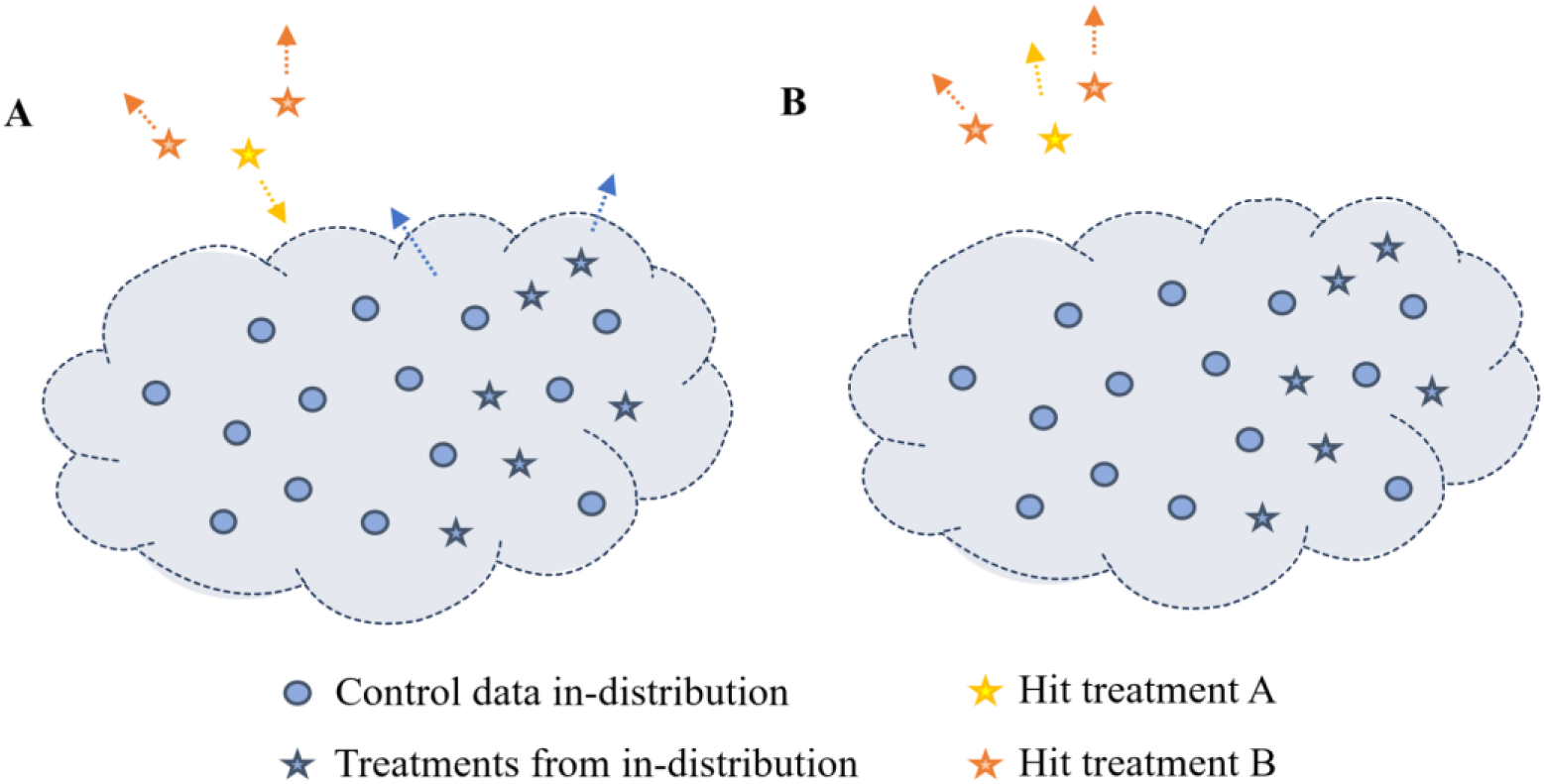
Illustration of potential caveats of using weakly supervised (e.g., using the treatment as a weak label) (A) versus anomaly-based (B) representations. (**A**) A treatment representation (yellow) is implicitly becoming more similar to the control because it was guided away from another treatment (orange). (**B**) Anomaly-based representations are guided away from the control.

**Supplementary Figure 6.**
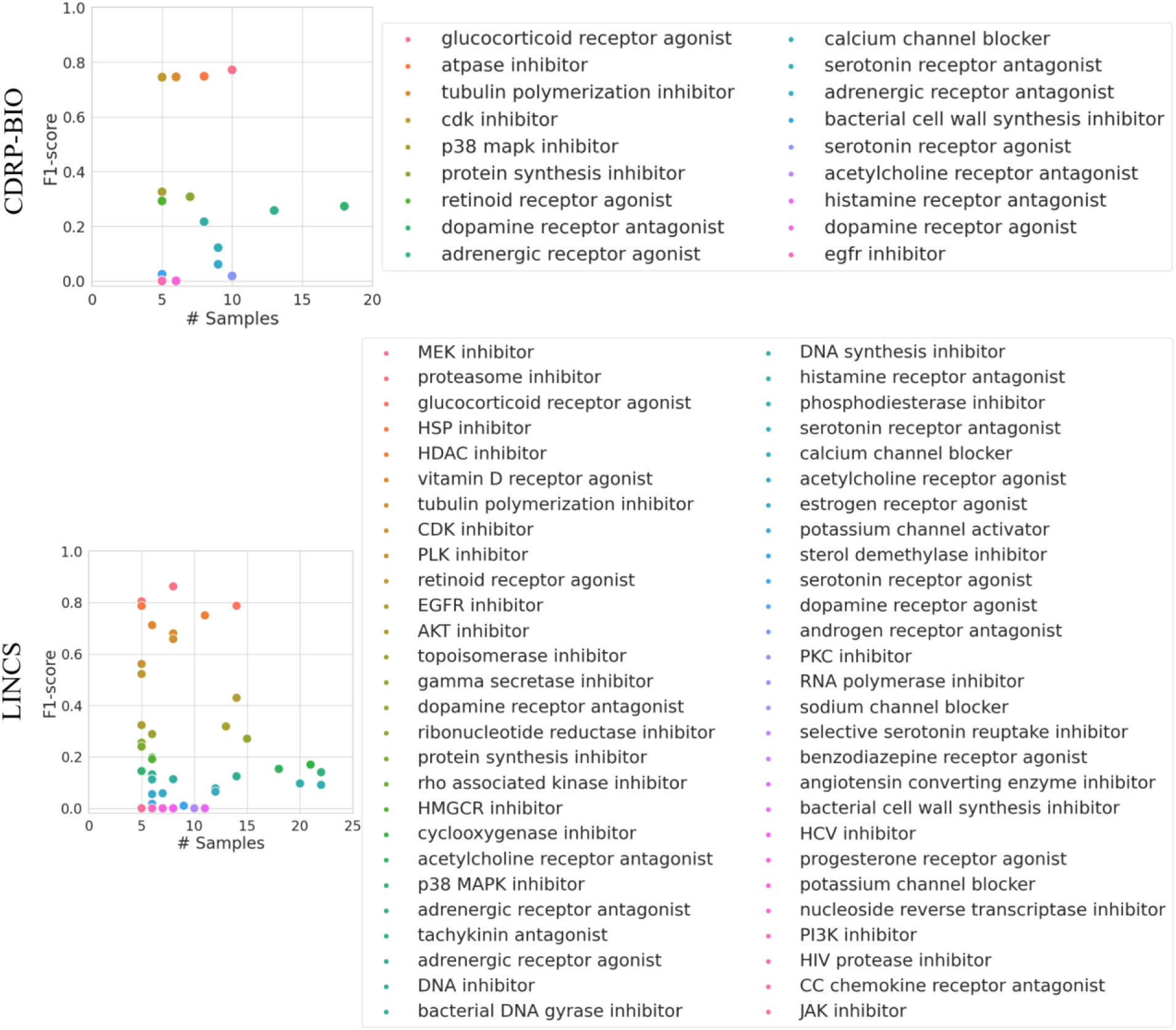
Scatter plot indicating the F1 score as a function of the number of treatments for each MoA in the CDRP-bio and LINCS dataset. The F1 scores were obtained with the better performing LR model trained using the anomaly-based representations.

## Funding and Acknowledgments

This research was supported by the Israel Science Foundation (ISF, grant No. 2516/21, to AZ), by the Israel Ministry of Science and Technology (MOST, to AZ), by the Wellcome Leap Delta Tissue program (to AZ), and by the Center for Open Bioimage Analysis (COBA) funded by the National Institute of General Medical Sciences P41 GM135019 (to EW). We thank Hillel Bar-Gera for his support and mentorship to AS.

## Author Contribution

AZ conceived the study. AS and NK developed the computational method and analyzed the data. AS, NK, SG, EW and AZ interpreted the results. AS and AZ drafted the manuscript. AZ acquired funding and mentored AS and NK. All authors edited the manuscript and approved its content.

